# STAMarker: Determining spatial domain-specific variable genes with saliency maps in deep learning

**DOI:** 10.1101/2022.11.07.515535

**Authors:** Chihao Zhang, Kangning Dong, Kazuyuki Aihara, Luonan Chen, Shihua Zhang

**Affiliations:** International Research Center for Neurointelligence, The University of Tokyo Institutes for Advanced Study, The University of Tokyo, Tokyo 113-0033, Japan; NCMIS, CEMS, RCSDS, Academy of Mathematics and Systems Science, Chinese Academy of Sciences, Beijing 100190, China; School of Mathematical Sciences/Sino-Danish College, University of Chinese Academy of Sciences, Beijing 100049, China; Key Laboratory of Systems Biology, Shanghai Institute of Biochemistry and Cell Biology, Center for Excellence in Molecular Cell Science, Chinese Academy of Sciences, Shanghai 200031, China; Key Laboratory of Systems Health Science of Zhejiang Province, School of Life Science, Hangzhou Institute for Advanced Study, University of Chinese Academy of Sciences, Hangzhou, 310024, China; School of Life Science and Technology, Shanghai Tech University, Shanghai 201210, China; Guangdong Institute of Intelligence Science and Technology, Hengqin, Zhuhai, Guangdong 519031, China

**Keywords:** spatial transcriptomics, spatial domain, spatially variable genes, deep learning, saliency map

## Abstract

Spatial transcriptomics characterizes gene expression profiles while retaining the information of the spatial context, providing an unprecedented opportunity to understand cellular systems. One of the essential tasks in such data analysis is to determine spatially variable genes (SVGs), which demonstrate spatial expression patterns. Existing methods only consider genes individually and fail to model the inter-dependence of genes. To this end, we present an analytic tool STAMarker for robustly determining spatial domain-specific SVGs with saliency maps in deep learning. STAMarker is a three-stage ensemble framework consisting of graph-attention autoencoders, multilayer perceptron (MLP) classifiers, and saliency map computation by the backpropagated gradient. We illustrate the effectiveness of STAMarker and compare it with three competing methods on four spatial transcriptomic data generated by various platforms. STAMarker considers all genes at once and is more robust when the dataset is very sparse. STAMarker could identify spatial domain-specific SVGs for characterizing spatial domains and enable in-depth analysis of the region of interest in the tissue section.

## Introduction

Knowing the relative spatial context of complex tissues or cell cultures is crucial to understanding complex biological systems^1^. The recent advances in spatial transcriptomic (ST) technologies have enabled gene expression profiling with spatial localization information. Such techniques (e.g., 10x Visium^2^, Slide-seq^3,4^, and Stereo-seq^5^) can profile the gene expressions corresponding to captured locations (referred to as spots or beads) at a resolution of several cells or even at a subcellular level, allowing us to discover spatially variable genes^6–8^, identify spatial domains^9–11^(i.e., regions with similar spatial expression patterns), deconvolve cell types of spots or beads^12,13^, and characterize spatial cell-to-cell interactions^14,15^.

One of the fundamental tasks in ST data analysis is to identify genes whose expressions display spatially varying patterns, simply referred to as spatially variable genes (SVGs). Common methods for identifying SVGs include trensceek^6^, SpatialDE^7^, SPARK^8^, SPARK-X^16^, and Hotspot^17^. The first three methods were designed based on the parametric framework. For example, SpatialDE fits a Gaussian process regression (GPR) model for each gene’s expression and finds whether the GPR model with spatial terms describes data better than that without using a log-likelihood ratio test. Fitting GPR models for large-scale ST data can be very time-consuming. To address this issue, SPARK-X adopts a non-parametric framework to test the dependence between each gene’s expression covariance and spatial covariance. Hotspot adopts a spatial autocorrelation metric (i.e., a modified Geary’s C statistics) to construct a test statistic to identify SVGs.

There are two main limitations of the existing methods. First, all the methods perform the hypothesis tests for each gene independently, ignoring the fact that the genes’ spatial expression patterns could be complementary to each other. Since ST data tend to be very sparse, performing hypothesis tests for genes individually may result in deteriorating performance. Second, the identified genes are not spatial domain-specific, hindering in-depth downstream analysis. For example, researchers may be interested in genes that display spatial patterns in one or several specific regions. However, none of the existing methods are directly applicable for such a purpose.

To this end, we propose an analytic tool STAMarker based on a three-stage ensemble framework consisting of graph-attention autoencoders, multilayer perceptron (MLP) classifiers, and saliency map computation by the backpropagated gradient to determine robust spatial domain-specific SVGs. Different from testing genes individually as the existing methods, STAMarker considers all genes at once by the backpropagated gradient (i.e., saliency map) and further identifies the spatial domain-specific SVGs. The intuition behind STAMarker is that genes contributing most to the tissue structures are potentially important to the corresponding spatial domains. The prominent advantage of STAMarker is its ability to identify spatial domain-specific SVGs, enabling deeper insight into specific regions. Extensive experiments and the comparison with the existing methods SpatialDE, SPARK-X and Hotspot on various ST data generated from different platforms (e.g., 10x Visium^2^, Slide-seq^3–5^, and Stereo-seq^5^) have shown its effectiveness.

## Results

### Overview of STAMarker

STAMarker is a three-stage framework that consists of an ensemble of graph attention autoencoders (STAGATE^10^), an ensemble of MLP classifiers and saliency map computation by the backpropagated gradient (**Fig. 1**). More specifically, after constructing the spatial neighbor network (SNN) based on the spots’ locations, STAMarker first trains multiple STAGATE graph-attention autoencoders, each of which learns the low-dimensional embeddings of the spots. The learned lowdimensional embeddings are used to identify spatial domains with various clustering algorithms, such as Louvain^18^ and mclust^19^. To obtain robust and unified spatial domains, STAMarker uses consensus clustering to aggregate the clustering results. Second, STAMarker trains multiple MLPs to model the relationship between the corresponding embeddings and the spatial domains. Third, to detect the SVGs, STAMarker stacks the encoders with the corresponding MLP and computes the saliency maps by backpropagation (see the “Saliency score” subsection of the Methods). STAMarker selects the SVGs in each spatial domain by their norms in the saliency maps.

**Figure 1.**
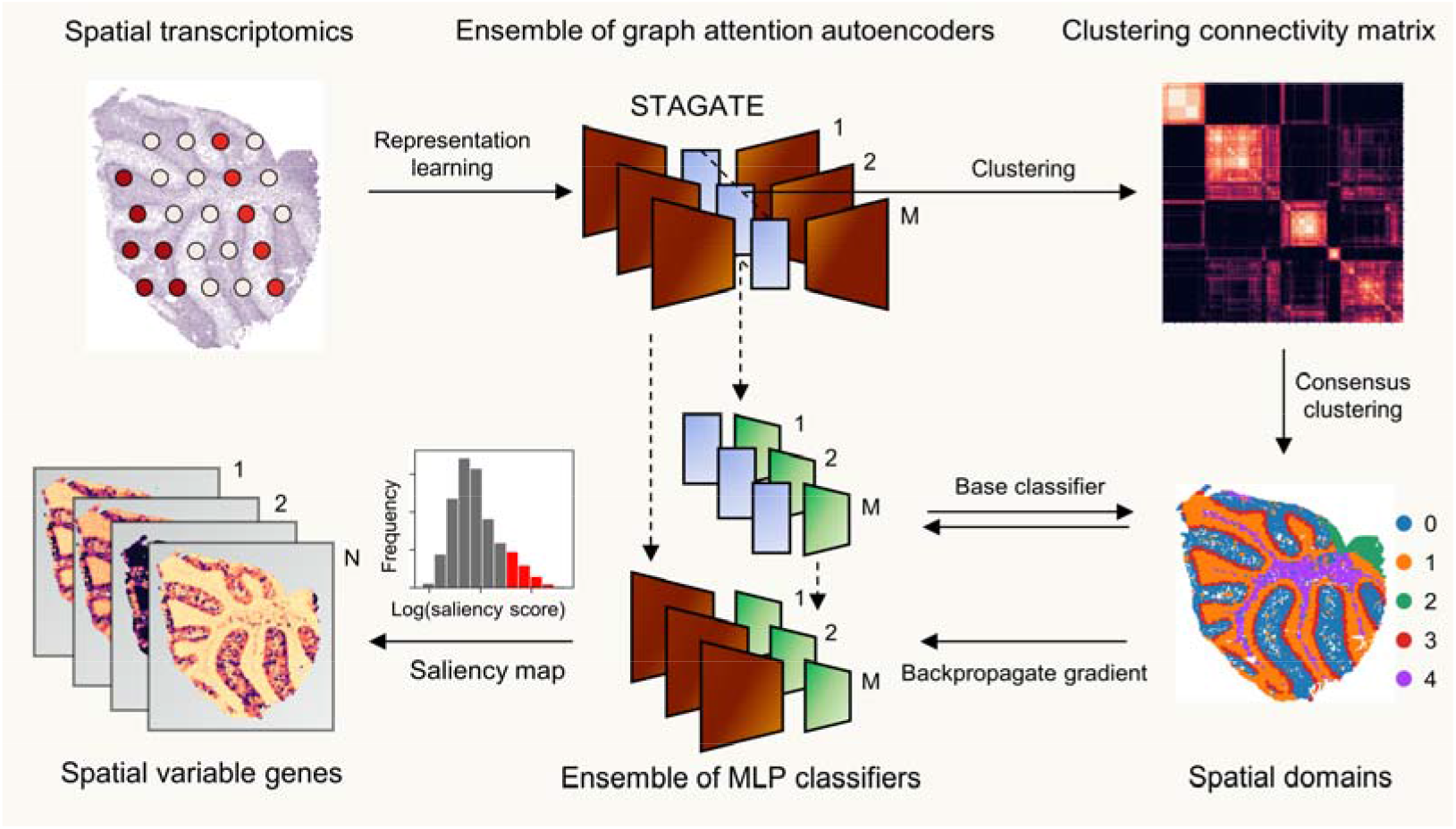
Overview of STAMarker. Given the spatial transcriptomics of a tissue section, STAMarker first trains an ensemble of graph attention auto-encoders that consists of *M* STAGATE models to learn the low-dimensional latent embeddings of spots, cluster them to obtain *M* grouping results, computes the clustering connectivity matrix and applies hierarchical clustering to obtain the spatial domains. STAMarker further models the relationships between the embeddings of the *M* auto-encoders and the spatial domains by training *M* base classifiers. At last, STAMarker computes the saliency map by first stacking the encoder and the corresponding classifier and then backpropagating the gradient to the input spatial transcriptomics matrix. STAMarker selects the domain-specific SVGs based on the genes’ saliency scores in each spatial domain.

### STAMarker robustly identifies the spatial domain-specific SVGs on the human dorsolateral prefrontal cortex dataset

We first applied STAMarker to the human dorsolateral prefrontal cortex (DLPFC) dataset profiled by the 10x Visium platform^20^. This dataset contains 12 sections that are manually annotated with the DLPFC layers and white matter (WM) based on the gene markers and morphological features (**Fig. 2A**). We considered the manual annotation as the ground truth and used the adjusted rand index (ARI) to evaluate the performance of spatial domain identification.

**Figure 2.**
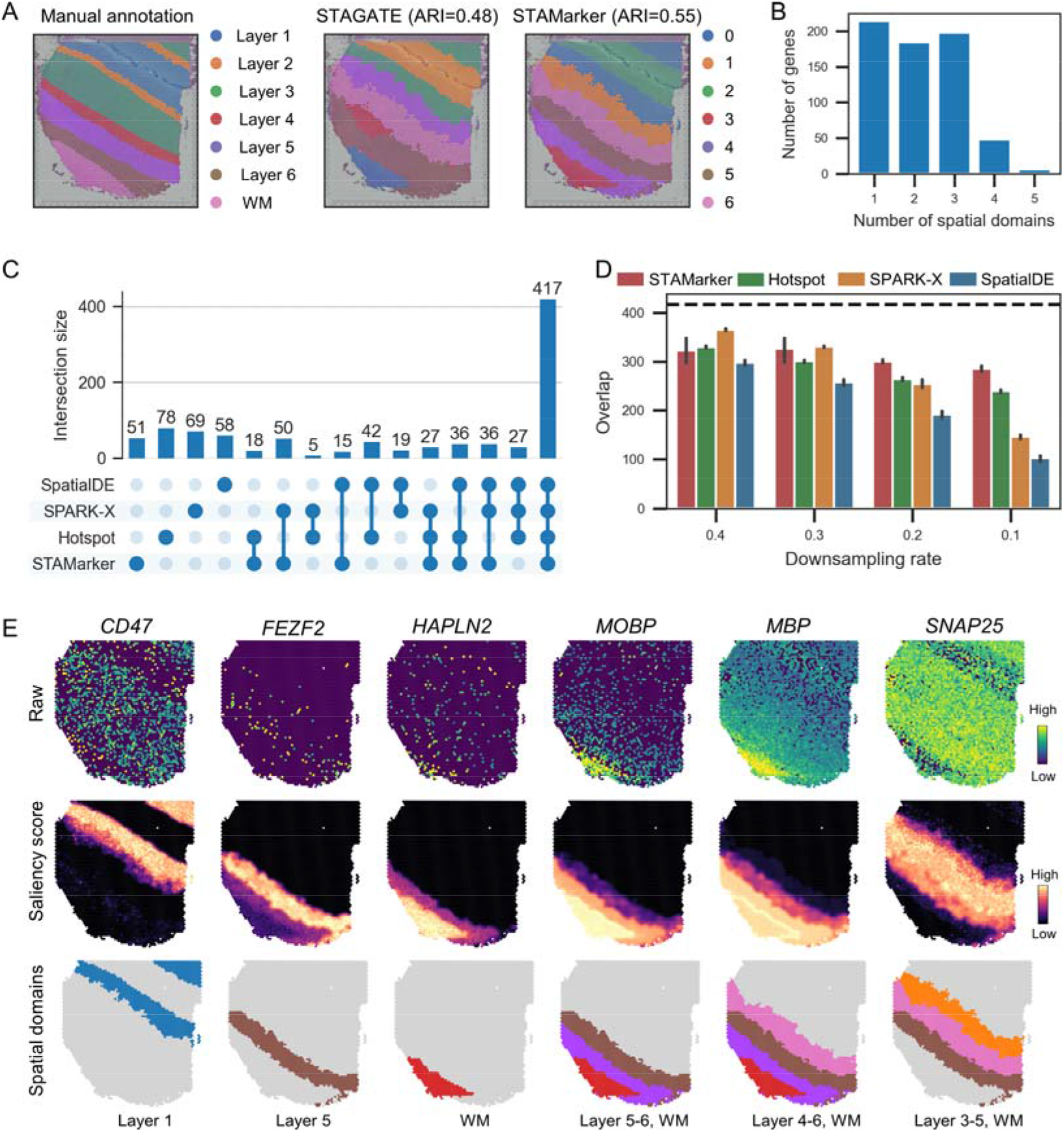
STAMarker robustly identifies the spatial domain-specific SVGs on the human dorsolateral prefrontal cortex (DLPFC) dataset. **A**, Manual annotation of cortical layers and white matter (WM) in the DLPFC section 151507, and the spatial domains identified by STAGATE and STAMarker respectively. **B**, Histogram of the number of spatial domains to which the SVGs identified by STAMarker belong. **C**, UpSet plot of the numbers of SVGs identified by SpatialDE, SPARK-X, Hotspot, and STAMarker. **D**, Comparison of the overlap number of the identified SVGs with the consensus ones by the four methods on the downsampled datasets. The error bars are computed based on five replicates. **E,** Visualization of the representative spatial domain-specific SVGs. From top to bottom, the raw counts, the saliency map (z-score transformation is applied), and the corresponding spatial domains of the SVGs identified by STAMarker.

Compared with only using an individual graph attention auto-encoder, STAMarker improved the robustness and identified the spatial domains more accurately. For example, STAGATE resulted in an undesirable spatial domain in the DLPFC section 151507 (the red region in **Fig. 2A**, middle panel). STAMarker improved the performance of ARI from 0.48 to 0.55 and identified the expected cortical layer structures better. Experimental results in the 12 DLPFC sections showed that STAMarker could consistently reduce the model variances of STAGATE across all sections and improve the performance of spatial domain identification in most of the sections (**Fig. S1B**). We set the number of auto-encoders *M* = 20 in all the following experiments.

STAMarker further identified the spatial domain-specific SVGs by saliency map. We note that a gene could be identified as an SVG for multiple spatial domains, which means that this gene is important for the determination of those spatial domains. For section 151507 of the DLPFC dataset, we found that 214 SVGs genes were specifically related to a unique spatial domain, and 382 SVGs were specifically with two or three spatial domains (**Fig. 2B**).

We compared STAMarker with three commonly used methods, i.e., SpatialDE^7^, SPARK-X^16^, and Hotspot^17^. Note that none of the compared methods can identify domain-specific SVGs. To facilitate direct comparison, we kept the number of the identified SVGs to be the same for all methods. The UpSet plot shows that the SVGs identified by the four methods have considerable overlap (417 out of 650 genes, referred to as consensus SVGs in the following), suggesting that STAMarker could discover the most SVGs identified by other methods (**Fig. 2C**). To further illustrate the robustness of STAMarker, we downsampled the counts of the gene expressions and computed the overlap number of the identified SVGs with the consensus ones (**Fig. 2D**). STAMarker is more robust than other methods when the gene expressions are sparse (the downsampling rate is less than 0.3) (**Fig. 2D**). The compared methods perform the hypothesis test for each independently, resulting in weak statistical power when the data are very sparse. STAMarker implicitly takes account of the gene expression interaction by considering all the genes at once with the backpropagation of the gradient, suggesting that it has a stronger power when the ST data are very sparse.

More importantly, STAMarker could determine spatial domain-specific SVGs that play important roles in DLPFC (**Fig. 2E**). We note that a gene that is highly expressed in a spatial domain does not necessarily have a higher saliency score in that region. For example, *CD47* was highly expressed in more than one layer, and it was identified as a layer 1-specific SVG by STAMarker but not by other methods. Note that *CD47* was known as the ligand of tyrosine phosphatases^21^, and was documented as an Alzheimer’s resilience factor^22^ which is related to the biological function of DLPFC. Moreover, STAMarker revealed *FEZF2* and *HAPLN2* as layer 5 and WM-specific genes respectively. Notably, *FEZF2* is a marker gene of deep layer excitatory neurons^23^, and *HAPLN2* plays an important role in the development of white matter^24^. Lastly, STAMarker also revealed some SVGs like *MOBP, MBP*, and *SNAP25* for multiple spatial domains, which have been reported as marker genes before^20^.

The saliency map can be used to cluster the spatial domain-specific SVGs into spatial modules (see the “**Identifying spatial domain-specific SVG modules**” subsection). The selected SVGs were clustered into seven clear modules (**Fig. S1C**) which correspond to the layers 1-6 and white matter, respectively (**Fig. S1D**). We also performed gene enrichment analysis for the identified SVGs by the four methods and found that the SVGs identified by STAMarker tended to be more enriched in GO cellular components (GO:CC) terms directly related to the nervous system, such as synapse and neuron projection (**Fig. S1E**). We observed consistent phenomena in other datasets (see Supplementary Notes section “**Comparison of the enrichment analysis of the identified SVGs by the four methods**” for detailed description).

### STAMarker enables fine-grained analysis on the mouse hippocampus dataset of Alzheimer’s disease

We applied STAMarker to the mouse hippocampus dataset of Alzheimer’s disease (AD)^25^, which was generated by Slide-seq V2 with a spatial resolution of 10μm. STAMarker could well characterize the tissue structures (**Fig. 3A**) including the important ones in the hippocampus, such as the arrow-like structure DG and the cord-like structure CA1. Strikingly, STAMarker successfully identified a spatial domain (domain 9) corresponding to the microglial cells (**Fig. 3B**), which were concentrated around amyloid plaques. This phenomenon is a prominent feature of AD^26,27^. STAMarker identified the SVGs of spatial domain 9, and many of them are known gene markers of microglial cell and risk genes of AD with significantly high expressions (**Fig. 3C**). For example, *P2ry12* was related to microglial motility and migration^28^; *Trem2* was known selectively expressed by microglia and related to a cell surface protein^27^. *Trem2* was also a well-known risk gene associated with AD, and its mutation increases the risk of AD around threefold^29,30^. *Hexb* and *Cx3rc1* were stably expressed microglia core genes^31^. However, among the competing methods, SpatialDE missed *Hexb* and *Fcrls*. SPARK-X only identified *Mef2c*, and Hotspot was the only method that could identify all the six known marker genes (shown in **Fig. 3C**). Compared with the competing spatial domain-agnostic methods, STAMarker enables a fine-grained analysis of the spatial domain of interest.

**Figure 3.**
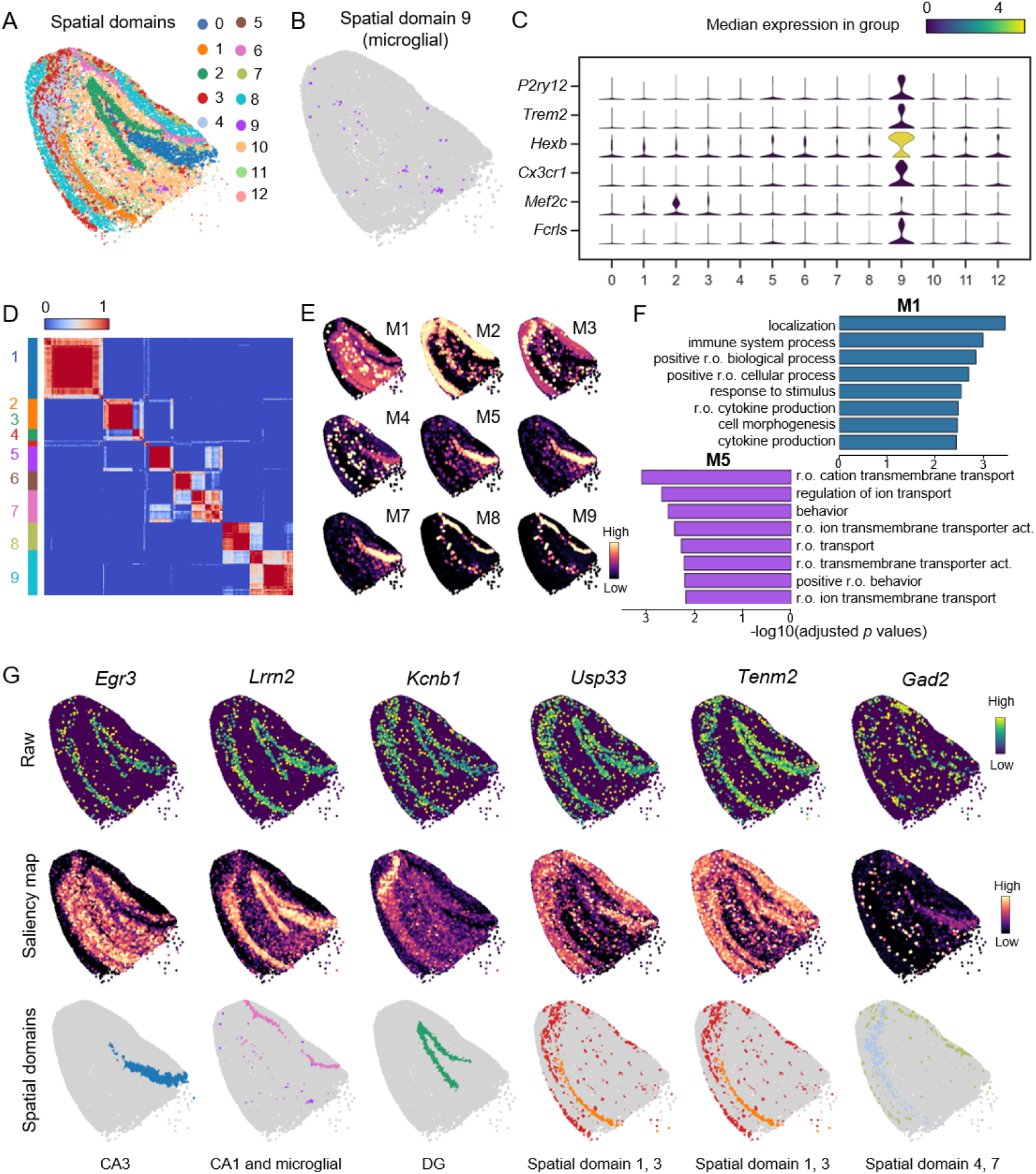
STAMarker identifies spatial domain-specific SVGs on the mouse hippocampus dataset. **A**, Spatial domains identified by STAMarker. The major subfields of the hippocampal formation, such as DG (spatial domain 2), CA1(spatial domain 6), and CA3 (spatial domain 0), were clearly shown. **B**, Spatial domain 9 corresponds to the microglial cells which are associated with the amyloid plaque of Alzheimer’s disease. **C**, Stacked violin plot showing the marker genes of microglial cells that were differentially expressed in spatial domain 9 but not others. **D**, Heatmap showing the nine clear gene modules that were clustered by the 154 spatial domain-specific SVGs. **E**, Visualization of the domain-specific gene modules by the first principal component of the saliency maps. **F**, Comparison of the top eight GO BP terms of M1 and M5. “r.o.” stands for “regulation of” to avoid clutter. **G**, Visualization of the representative spatial domain-specific SVGs. From top to bottom, the raw counts, the saliency map, and the corresponding spatial domains of SVGs.

In total, STAMarker determined 797 SVGs that had considerable overlap with the ones identified by other methods and many of the STAMarker-identified SVGs are domain-specific (**Fig. S2A and B**). These SVGs detected by STAMarker were enriched in a total of 1125 GO terms and 47 KEGG at an FDR of 5%. The directly related GO terms such as synapse (GO:0045202), nervous system development (GO:0007399), and neuron projection (GO:0043005) were significantly enriched and comparable to the competing methods (**Fig. S2C**).

We further used the saliency map to cluster SVGs belonging to less than two spatial domains (referred to as domain-specific SVGs for simplicity) into nine gene modules (**Fig. 3D**, 312 SVGs in total). Gene module M1 (74 genes) was highlighted in the microglial cells and was specifically enriched in many GO terms related to the immune response terms, such as immune system process, response to stimulus, and regulation of cytokine production (**Fig. 3F**). As a comparison, the enriched GO terms of gene module M5 were mainly related to transmembrane transport and no immune-related GO terms were detected. The representative genes of gene module M1 include *Egr3, Lrrn2, and Kcnb1* (**Fig. 3G**). *Egr3* was CA3-specific and only identified by STAMarker. *Egr3* was known as a master regulator of differentially expressed genes in AD^32^. *Lrrn2* corresponded to microglial and CA1. *Kcnb1* was a DG-specific one, indicating that it is important for distinguishing DG from other spatial domains. *Kcnb1* was also associated with aging and cognitive impairment^33^. Gene module M2 (37 genes) was mainly highlighted in the exterior region with two representative genes *Usp33* and *Tenm2*. This module was significantly enriched in axon growth(GO:0007409, *P* = 3.55 × 10^-26^) and axon guidance (GO:0007411, *P* = 2.74 × 10^-10^). Gene module M5 (30 genes) was enriched in many GO terms related to neurons such as regulation of neurotransmitter levels (GO:0001505, *P* = 5.78 × 10^-23^), and presynapse (GO:0098793, *P* = 2.85 × 10^-57^). Its representative gene *Gad2* was a spatially variable one related to spatial domains 4 and 7 (**Fig. 3G**).

### STAMarker reveals the domain-specific variable genes on the mouse olfactory bulb dataset

We applied STAMarker to the mouse olfactory bulb dataset^5^ profiled by the Stereo-seq platform to identify the laminar organization and the corresponding SVGs. STAMarker could well decipher the eight spatial domains with clear laminar structures (**Fig. 4A**), consisting of rostral migratory stream (RMS), ependymal cell zone (ECZ), granule cell layer (GCL), internal plexiform layer (IPL), mitral layer (MT), glomerular layer (GL) and olfactory nerve layer (ONL), annotated according to the Allen Brain Atlas^34^.

**Figure 4.**
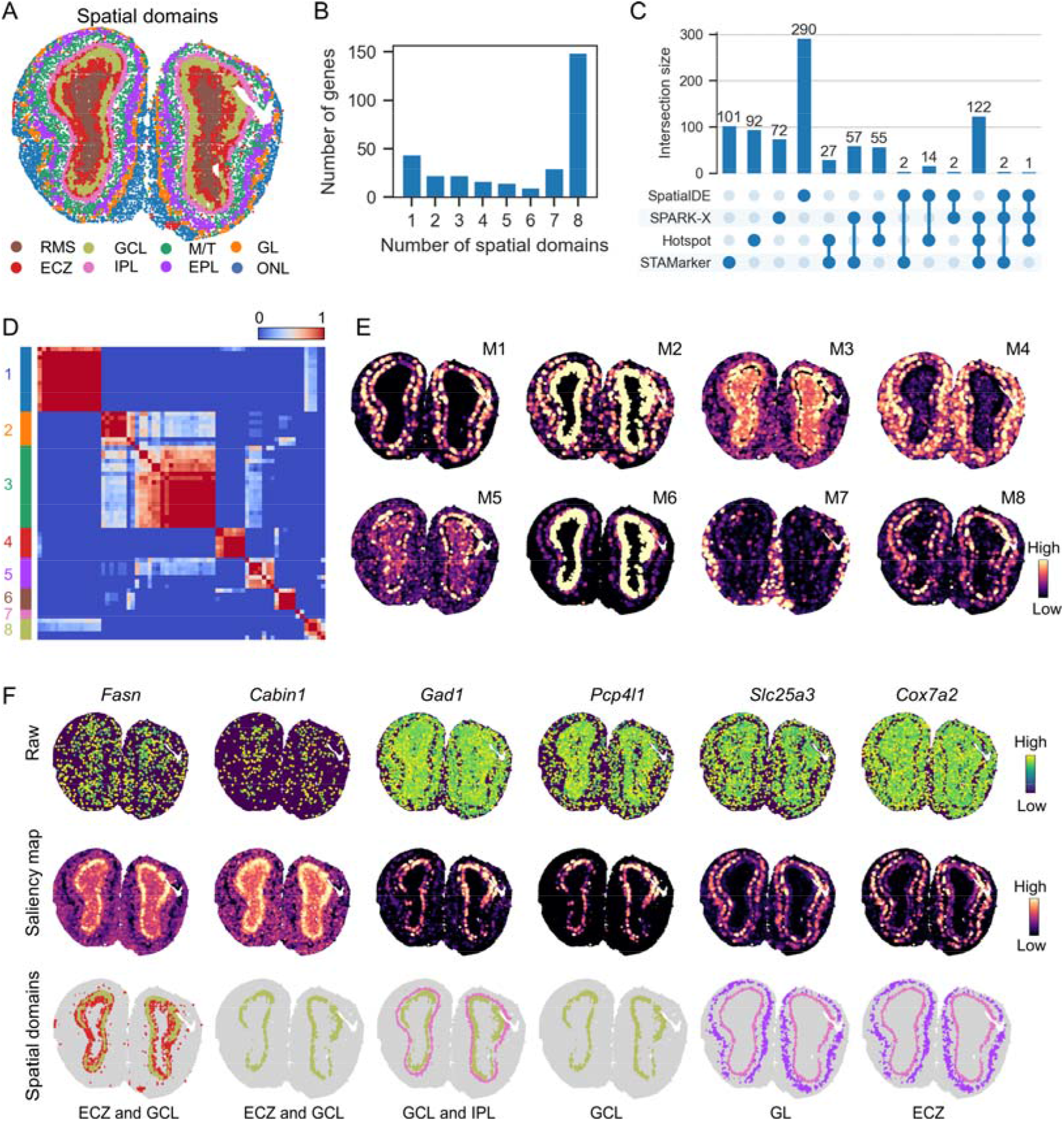
STAMarker reveals the domain-specific SVGs on the mouse olfactory bulb dataset. **A,** Spatial domains identified by STAMarker. The laminar organization of the mouse olfactory bulb is clearly shown. The identified spatial domains were annotated by the Allen Reference Atlas. **B**, Histogram of the number of spatial domains to which the SVGs identified by STAMarker belong. **C**, UpSet plot of the numbers of SVGs identified by SpatialDE, SPARK-X, Hotspot, and STAMarker. STAMarker identified 311 SVGs. **D**, Heatmap showing the eight modules that were clustered by the 67 domain-specific SVGs. **E**, Visualization of the domain-specific gene modules by the first principal component of the saliency maps. **F**, Visualization of the representative spatial domain-specific SVGs.

STAMarker determined 311 SVGs, and most of them belong to more than three spatial domains (**Fig. 4B**), implying that the eight laminar organizations share similar gene expression patterns. The shared SVGs of the four methods is only one gene (**Fig. 4C**). The SVGs identified by STAMarker were enriched in more GO terms and KEGG pathways, while those of SpatialDE were enriched in much fewer terms at an FDR of 5% (**Fig. S3**) (STAMarker: 451 GO terms and 42 KEGG pathways; Hotspot: 349 GO terms, 27 KEGG pathways; SPARK-X: 377 GO terms, 34 KEGG pathways; SpatialDE: 15 GO terms, 2 KEGG pathways). Many of the STAMarker-identified GO terms and KEGG pathways directly related to the synapse organization and the functions of the olfactory bulb, and tended to be more significant than those of the other methods. For example, the SVGs identified by STAMarker were enriched in the circadian entrainment pathway (KEGG 04713, *P* = 4.13 × 10^-12^) and were more significant than those by the compared methods (Hotspot *P* = 3.05 × 10^-7^; SPARK-X *p* = 10^-6^; SpatialDE is not significant).

The spatial domain-specific SVGs (67 genes) were clustered into eight gene modules (**Fig. 4D**), showing clear laminar organization with distinct correspondence to the morphological layers. For example, gene module M1 consisted of spatial domain-specific SVGs of the ECZ and GCL layers (**Fig. 4E**). Its two representative genes *Fasn* and *Cabin1* were, respectively, related to the endothelial cell differentiation (GO 0045446, *P* = 6.60 × 10^-3^) and over-expressed in olfactory epithelium^35^ (**Fig. 4F**), which was only identified by STAMarker. Gene module M3 highlighted the interior layers of the olfactory bulb with two typical genes: *Gad1* was highlighted in both GCL and IPL and played an important role in the olfactory bulb interneurons^36^ and *Pcp4l1* was known as a marker gene of the GCL^37^. Gene module M4 mainly corresponded to the IPL and EPL spatial domains with two typical genes *Slc25a3* and *Cox7a2*, and was enriched in the organelle inner membrane (GO 0019866, *P* = 6.91 × 10^-8^).

### STAMarker uncovers the spatial domain-specific SVGs on the mouse cerebellum dataset

We illustrated the effectiveness of STAMarker on a mouse cerebellum dataset generated by Slide-seq V2 platform^38^. STAMarker clearly identified the tissue structures of the cerebellum, i.e., molecular layer (MOL), Purkinje layer (PL), white matter (WM) and granule cell layer (GCL), and cerebellar nucleus (CN) (**Fig. 5A**). STAMarker and other three methods determined 508 SVGs respectively and the SVGs identified by SpatialDE were quite different from those of other methods (**Fig. S4**). The SVGs identified by STAMarker were enriched in 640 GO terms and 30 KEGG pathways, while the SVGs identified by SPARK-X were enriched in 578 GO terms and 32 pathways (for comparison, Hotspot: 550 GO terms and 24 KEGG pathways; SpatialDE: 170 GO terms and no enriched KEGG pathways were identified) (**Fig. S4C**). STAMarker identified the GO terms that are directly related the cerebellum development (GO 0021549, *P* = 3.59 × 10^-4^) and morphogenesis (GO 0021587, *P* = 4.83 × 10^-4^), while Hotspot is the only method that identified the two GO terms with less overlap size (GO 0021549, *P* = 5.67 × 10^-3^; GO 0021587, *P* = 0.160). One of the prominent advantages of STAMarker is the ability to provide spatial domain-specific SVGs, enabling more fine-grained analysis. Moreover, the spatial domain-specific enrichment analysis characterized the functions of each spatial domain. For example, the Purkinje-related GO terms were only enriched in MOL, PL, and GCL respectively (**Fig. 5B**), which is consistent with the histology of Purkinje cells.

**Figure 5.**
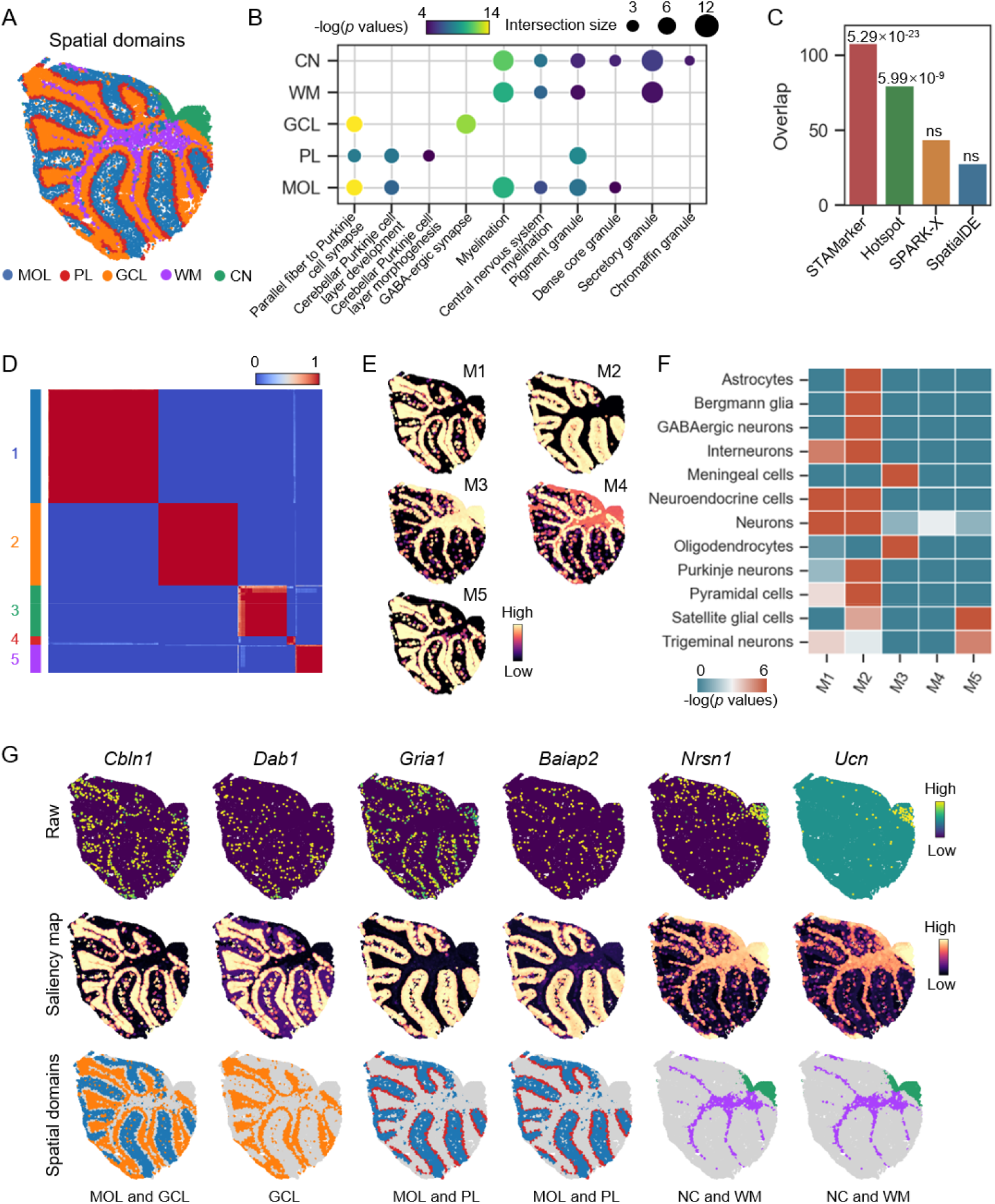
STAMarker uncovers spatial domain-specific SVGs on the mouse cerebellum data. **A,** Spatial domains identified by STAMarker. The identified spatial domains were annotated by the Allen Reference Atlas. **B**, GO enrichment analysis of the SVGs in the named spatial domains. The selected enriched GO terms show distinct significance levels. **C**, Bar plot displaying the overlap of the identified SVGs with reference gene list (261 genes, obtained from the Harmonizome database). The *p* values of the hypergeometric test were shown above the bars (ns indicates not significant, i.e., *p* values > 0.05). **D**, Heatmap showing the five gene modules that were clustered by the 369 domain-specific SVGs. **E**, Visualization of the domain-specific gene modules by the first principal component of the saliency maps. **F**, Enrichment analysis of the identified spatial modules with diverse cell types related to the cerebellum. **G**, Visualization of the representative spatial domain-specific SVGs.

Moreover, the SVGs determined by STAMarker could cover more mouse cerebellum marker genes from the Harmonizome database^39^ (i.e., 108 out of the 261 genes) with the hypergeometric test *P* = 5.29 × 10^-23^ compared to the other three methods (**Fig. 5C**). Hotspot identified the second most 80 genes with the hypergeometric test *P* = 5.99 × 10^-9^, while the overlaps of the identified SVGs by SPARK-X and SpatialDE were merely below 50 without statistical significance.

The spatial domain-specific SVGs were clearly clustered into five gene modules based on the saliency maps (**Fig. 5D**) with clear spatial domain patterns (**Fig. 5E**). The five modules were enriched in distinct cell types (**Fig. 5F**) based on the downloaded marker genes of cerebellum-related cell types from PanglaoDB^40^. For example, gene module M1 corresponds to GCL and MOL with two representative genes *Cbln1* and *Dab1*. The two genes were strongly over-expressed in the corresponding spatial domains and were enriched in cerebellum development (GO 0021549, *P* = 4.13 × 10^-8^). *Cbln1* was essential for synaptic plasticity and integrity in the cerebellum^41^. Gene module M2 mainly consists of PL and MOL and the Purkinje neurons were only enriched in this module. Two example genes of M2 include *Gria1* and *Baiap2* belonging to the enriched GO term regulation of synaptic plasticity (GO 0048167, *P* = 3.49 × 10^-9^). In particular, *Baiap2* was only identified by STAMarker. Gene module M5 was related to the CN and WM spatial domains with two highly expressed genes *Nrsn1* and *Ucn* in the CN region. Moreover, both of them were in the enriched GO term distal axon (GO 0150034, *P* = 3.88 × 10^-19^). In summary, STAMarker can not only determine SVGs that are enriched in the most GO terms and KEGG pathways that are directly related to the tissue biological functions compared to other methods, but also provides the spatial domains corresponding to SVGs.

## Discussion

Determining SVGs is the very first step towards understanding the complex biological functions of complex tissues and cell cultures through ST data. Here, we develop a robust and effective spatial domain-specific SVGs identification method STAMarker. STAMarker is a three-stage analytic method inspired by the saliency map in deep learning. STAMarker conceptually considers genes that contribute most to the determination of the spatial domains as SVGs. Different from the competing methods that consider genes independently, STAMarker considers all genes simultaneously, making it possible to exploit the complementary information across genes. Another outstanding advantage of STAMarker is its ability to identify spatial domain-specific SVGs while the competing methods are not directly applicable.

STAMarker could robustly identify SVGs even when the data are downsampled to a very sparse level. Experimental results on various datasets consistently showed that SVGs identified by STAMarker tended to be more significant in related GO terms and KEGG pathways. Moreover, the spatial domain-specific SVGs can be organized into gene modules corresponding to spatial domain patterns. The domain-specific SVGs identified by STAMarker could also enable researchers to investigate spatial domains of interest at a finer scale.

Despite that STAMarker considers all genes simultaneously and measures genes’ importance by their contributions to the classification, it is still unclear how to evaluate the importance of a set of genes, which are expected to be solved in the future. Lastly, we expect that this kind of analytic tools can be extended to other spatial omics like spatial metabolomics^42^.

## Materials and Methods

### Data description

We applied STAMarker and the three compared methods to ST datasets generated by various techniques, including 10x Visium, Slide-seqV2, and Stereo-seq (see **Supplementary Table S1** for details).

### Data preprocessing

We followed the data preprocessing procedure in STAGATE^10^. We first removed spots outside of the main tissue area. We then used the pipeline provided by the SCANPY package to log-transform the raw gene expression and normalize it according to the library size. For all datasets, we selected the top 3,000 highly variable genes as the inputs of STAMarker.

### Ensemble of graph attention auto-encoders

We used an ensemble of graph attention auto-encoders to robustly identify spatial domains. We adopted STAGATE as the base auto-encoder for its superior performance in spatial domain identification. Specifically, STAGATE consists of three parts: encoder *f_enc_*, decoder *f_dec_* and the attention layer. STAGATE first constructs a spatial neighbor network (SNN) where neighboring spots have edges. The encoder *f_enc_* transforms the gene expression of a spot into a *d*-dimensional embedding by aggregating information from the neighboring spots in SNN. Let’s denote the gene expression matrix of *n* spots and *p* genes by *X* ∈ *R*^*n*×*p*^. The encoder is defined as

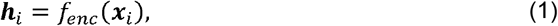

where *x_i_* ∈ *R^p^* indicates the gene expression profile of spot *i*. and *h_i_* ∈ *R^d^* is the corresponding latent embedding. Then the decoder *f_dec_* reverses back the latent embedding *h_i_* into the reconstructed gene expression profile, i.e., 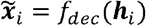. The attention layers coupled with the encoder and decoder adaptively learn the edge weights of SNN.

Despite that STAGATE has shown its effectiveness on various ST datasets, an inherent challenge of deep learning methods is that the weight initialization might have a significant impact on the results. To alleviate this issue, we constructed an ensemble of graph attention auto-encoders. We trained *M* STAGATE models with different random weight initialization and applied clustering algorithms to the learned low-dimensional embeddings of each model. Specifically, we used the mclust clustering algorithm when the number of labels is known; otherwise, we employed the Louvain algorithm. As a result, we obtained *M* clustering results for each STAGATE model, denoted by *Y*^(1)^, *Y*^(2)^,…, *Y*^(*M*)^, respectively.

### Consensus clustering

We used consensus clustering to obtain the spatial domains. Given the clustering results *Y*^(1)^, *Y*^(2)^,…, *Y*^(*M*)^, we first computed the clustering connectivity matrix *C* such that

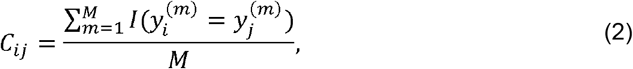

where *l*(·) is the indicator function, and 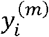 and 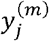 denote the cluster labels of spot *i*. and *j* in *Y*^(*m*)^ respectively. The clustering connectivity matrix *C* is a symmetric matrix where *C_ij_* indicates the empirical probability of spots *i*. and *j* that belong to the same cluster. Finally, we applied hierarchical clustering to *C* and obtained the spatial domain labels *Y*^*^. The number of clusters *K* is simply set as the same as the mode of the number of clusters of the *M* clustering results.

### Ensemble of MLP classifiers

We used an ensemble of two-layer MLP classifiers denoted by *f_MLP_* to model the relationship between the latent embeddings and the spatial domain labels *Y*^*^. The low-dimensional latent embeddings of the *M* auto-encoders are used as the inputs:

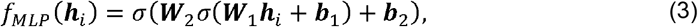

where *W*_1_ ∈ *R*^*d*×*d*^ and *W*_2_ ∈ *R*^*d*×*K*^ are the weight matrices, ***b***_1_ ∈ *R^d^*, ***b***_2_ ∈ *R^K^* are the bias, and *σ* is a nonlinear activation function (we used ReLU here). The MLP classifier is trained by minimizing the cross-entropy between the spatial domain labels *Y*^*^ and the softmax of the output of *f_MLP_*. As a result, there are *M* MLP classifiers corresponding to the *M* auto-encoders.

### Spatial domain-specific saliency map

We used the saliency map to measure the contribution of genes to the spatial domain classification. The idea is inspired by the saliency map in computer vision^43–45^, which is an important concept to explain the contribution of pixels to image classification. Specifically, given the *i*-th spot’s gene expression *x_i_*, we denoted the output of the last layer of the MLP classifier as *z_i_*

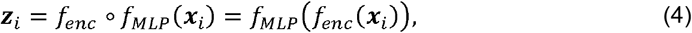

where ***z**_i_* ∈ *R^K^*. The MLP classifier will predict *x_i_* as the *k_max_*-th spatial domain where *k_max_* is the index of the largest value of *z_i_* (denoted by *z_k,k_max__*). Note that *z_i,k_max__* is a scalar function of *x_i_*. To measure the genes’ contribution to the prediction of spatial domains, we computed the gradient of *z_i,k_max__* with respect to *x_i_* by backpropagation

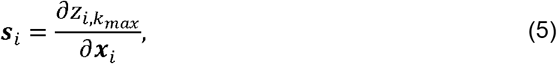

where *s_i_* ∈ *R^p^* is the same size of *x_i_*. We denoted the saliency map of the *m*-th auto-encoder and the corresponding MLP classifier across all spots as *S*^(*m*)^ ∈ *R*^*n*×*p*^, where the *i*-th column of is computed by (5). We used the norm of the gene *j*’s gradients across the spots belonging to spatial domain *k*. Specifically, the spatial domain-specific saliency score of gene *j* for spatial domain *k* is defined as follows:

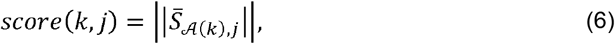

where 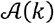 indicates the set of indices of spots belonging to spatial domain *k*, i.e., 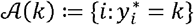.

### Identifying spatial domain-specific SVGs

We used the norm of the saliency maps to identify the spatial domain-specific SVGs. Given the spatial domain *k*, we first applied log transformation to the saliency scores of all genes. To adaptively select SVGs, we estimated the mean 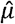 and standard deviation 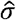 of the log-transformed scores of all genes, and selected genes whose scores are greater than 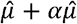, where *α* is a user-defined parameter to control the number of selected SVGs. Larger *α* results in fewer identified SVGs. We typically set *α* = 1.5.

### Identifying spatial domain-specific SVG modules

We constructed the spatial domain-specific SVG modules by the saliency score matrix. First, we selected genes that are SVGs in less than two spatial domains. Then we evaluated the pair-wise gene correlation by the saliency matrix, i.e., the Pearson correlation between 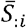 and 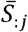. We then clustered the resulting affinity matrix into *K* clusters. We applied PCA to the saliency score matrix of the gene modules and visualized the gene modules by the first principal component.

### Gene enrichment analysis

We used the g:pforfie^46^ API provided by SCANPY^47^ to perform gene enrichment analysis.

## Data availability

All data used in this paper are available in raw from the corresponding papers’ authors. Specifically, the DLPFC dataset generated by 10x Visium is available at http://spatial.libd.org/spatialLIBD. The hippocampus dataset of the J20 mouse model generated by Slide-seq V2 is accessible at https://singlecell.broadinstitute.org/single_cell/study/SCP1663/cell-type-specific-inference-of-differential-expression-in-spatial-transcriptomics. The processed Stereo-seq data for mouse olfactory bulb tissue is available at https://github.com/JinmiaoChenLab/SEDR_analyses. The processed Slide-seq data for mouse cerebellum is available at https://singlecell.broadinstitute.org/single_cell/study/SCP354/slide-seq-study.

## Code availability

The STAMarker algorithm is implemented with Python and is available at http://github.com/zhanglabtools/STAMarker.

## Acknowledgments

This work was supported by the National Key Research and Development Program of China [No. 2019YFA0709501], JST Moonshot R&D Grant [No. JPMJMS2021], AMED [No. JP22dm0307009], JSPS KAKENHI [No. JP20H05921], Institute of AI and Beyond at the University of Tokyo, the Intelligent Mobility Society Design Social Cooperation Program at the University of Tokyo, the Strategic Priority Research Program of the Chinese Academy of Sciences (CAS) [Nos. XDA16021400, XDPB17], the National Natural Science Foundation of China [Nos. 12126605, 61621003], the Key-Area Research and Development of Guangdong Province [No. 2020B1111190001], and the CAS Project for Young Scientists in Basic Research [No. YSBR-034].

## Author contributions

S.Z. conceived and supervised the project. C.Z. developed and implemented the STAMarker package. C.Z., K.D., A.K., L.C., and S.Z. validated the methods and wrote the manuscript. All authors read and approved the final manuscript.

