## Supplementary Material for "STAMarker: Determining spatial domain-specific variable genes with saliency maps in deep learning"

**Supplementary Notes**

**Datasets**

We applied STAMarker to various datasets generated by different ST platforms to illustrate its effectiveness in this study (**Table S1**). Specifically, we applied STAMarker to the human dorsolateral prefrontal cortex (DLPFC) dataset generated by 10X Visium^1^, the mouse hippocampus dataset^2^ and cerebellum dataset^3^ generated by Slide-seq V2, and the mouse olfactory bulb dataset^4^ generated by Stereo-seq.

Specifically, we used section 151507 of the 10x Visium DLPFC dataset in the main text. It contains 4226 spots. We also applied STAMarker to the 12 sections to evaluate the performance of STAMarker (**Figure S1**). Compared with the 10x Visium platform which profiles genes at a resolution of ~55μm, Slide-seqV2 has a higher resolution but less sequence depth. In the mouse hippocampus dataset, we used the cropped data of the first replicate, which consists of 15092 spots. The mouse cerebellum dataset we used consists of 32701 spots. The Stereo-seq platform uses DNA nanoball-pattered array chips to achieve a subcellular spatial resolution. The mouse olfactory bulb dataset used here was processed into a cellular-level resolution (~14μm). As a consequence, it contains 19109 spots.

**Comparison with competing methods**

We compared STAMarker with three commonly used methods for spatially variable gene identification, including SpaitalDE^5^, SPARK-X^6^, and Hotspot^7^. To facilitate the comparison, the data preprocessing procedure is the same as described in the main text. We retained the top 3000 highly variable genes (HVGs) as the inputs of all the compared methods and followed the authors’ instructions for detailed implementation.

- **SpatialDE:** The raw counts of the selected top 3000 HVGs were normalized by *NaiveDE.stabilize* and the log-transformed data were regressed with respect to locations by *NaiveDE.regress_out*. Finally, the spatially variable genes were identified using *SpatialDE.run* with default parameters.
- **Hotspot:** The raw counts of the selected top 3000 HVGs were fitted by hotspot.Hotspot with the *danb* model. The KNN graph was created by *create_knn_graph* where the *n_neighbors* is set by 30. Finally, the autocorrelations were computed by *compute_autocorrelations()*.
- **SPARK-X:** The raw counts of the selected top 3000 HVGs and the locations were fitted by *sparkx* with the option set as “*mixture”.* There were eleven nonparametric models with different kernel functions were fitted. Finally, the spatially variable genes were selected by the combined *p* values.

**Comparison of the enrichment of the identified SVGs**

We performed gene enrichment analysis of the SVGs identified by the four methods. Specifically, we used the function *scanpy.queries.enrich* in SCANPY^8^ which provides API of g:profile^9^ to perform enrichment analysis. The FDR was set by 0.05 in all experiments. We plotted the *p* values of GO terms enriched with SVGs identified by STAMarker versus that of the compared methods on the four datasets (**Figure S1E**, **Figure S2C**, **Figure S3,** and **Figure S4C**). We annotated some of the GO terms in each plot.

In general, the SVGs identified by STAMarker tend to be more enriched in GO terms relevant to the tissues. In the DLPFC dataset, the relevant GO terms such as synapse, cell junction, somatodendritic compartment, and neuron projection are consistently above the diagonal line (**Figure S1E**), indicating that the SVGs identified by STAMarker are more significant than those of the compared methods. In the mouse hippocampus dataset, relevant GO terms including nervous system development, synapse, neuron projection, and others are more significant or comparable to the compared methods (**Figure S2C**). The result in the mouse olfactory data is consistent (**Figure S3**). The relevant GO terms including synapse, cell junction, and somatodendritic compartment are more significant (above the diagonal), while GO terms that are less specific to the tissues (e.g., endomembrane system, anatomical structure development, left panel; cytoplasm and membrane protein complex, middle panel; HAUS complex, right panel) are below the diagonal. In addition, the SVGs identified by SpatialDE are enriched in much fewer GO terms than the other three methods (**Figure S3**, right panel). In the mouse cerebellum data, the result is also consistent (**Figure S4C**); the less specific GO terms (e.g., intracellular anatomical structure) are below the diagonal, and the relevant GO terms (neuron projection and synapse) are above the diagonal.

**Supplementary Figures**

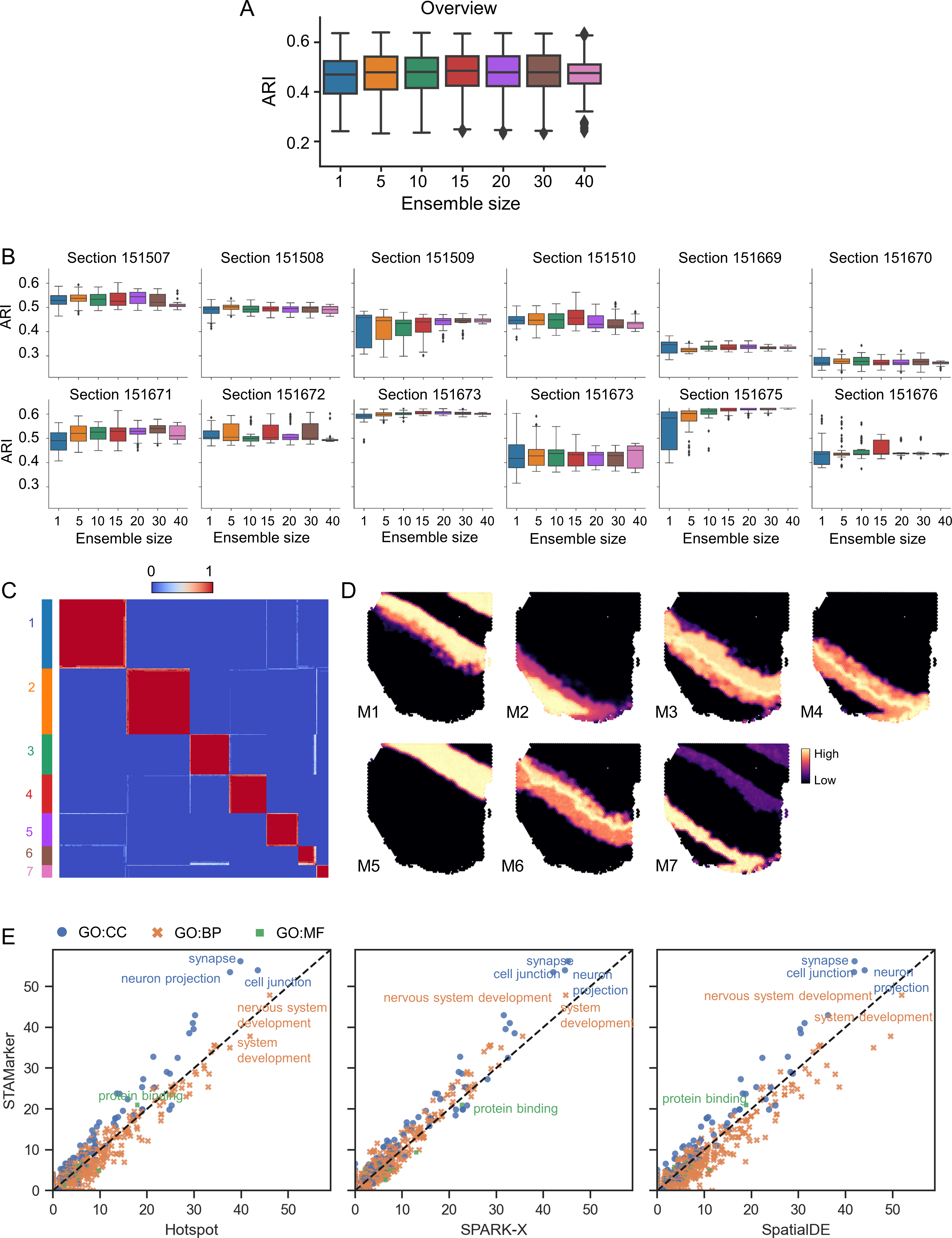

**Figure S1.** **Additional results on the DLPFC dataset**. **A** and **B**, Overall and individual performance of the spatial domains identified by STAMarker on the 12 sections with different numbers of autoencoders. **C**, Heatmap showing the seven clear gene modules clustered by the 154 spatial domain-specific SVGs. **D**, Domain-specific gene modules were visualized by the first principal component of the saliency scores. **E**, Comparison of enrichment significance –log(*p* values) of GO terms enriched with SVGs identified by STAMarker and three compared methods.

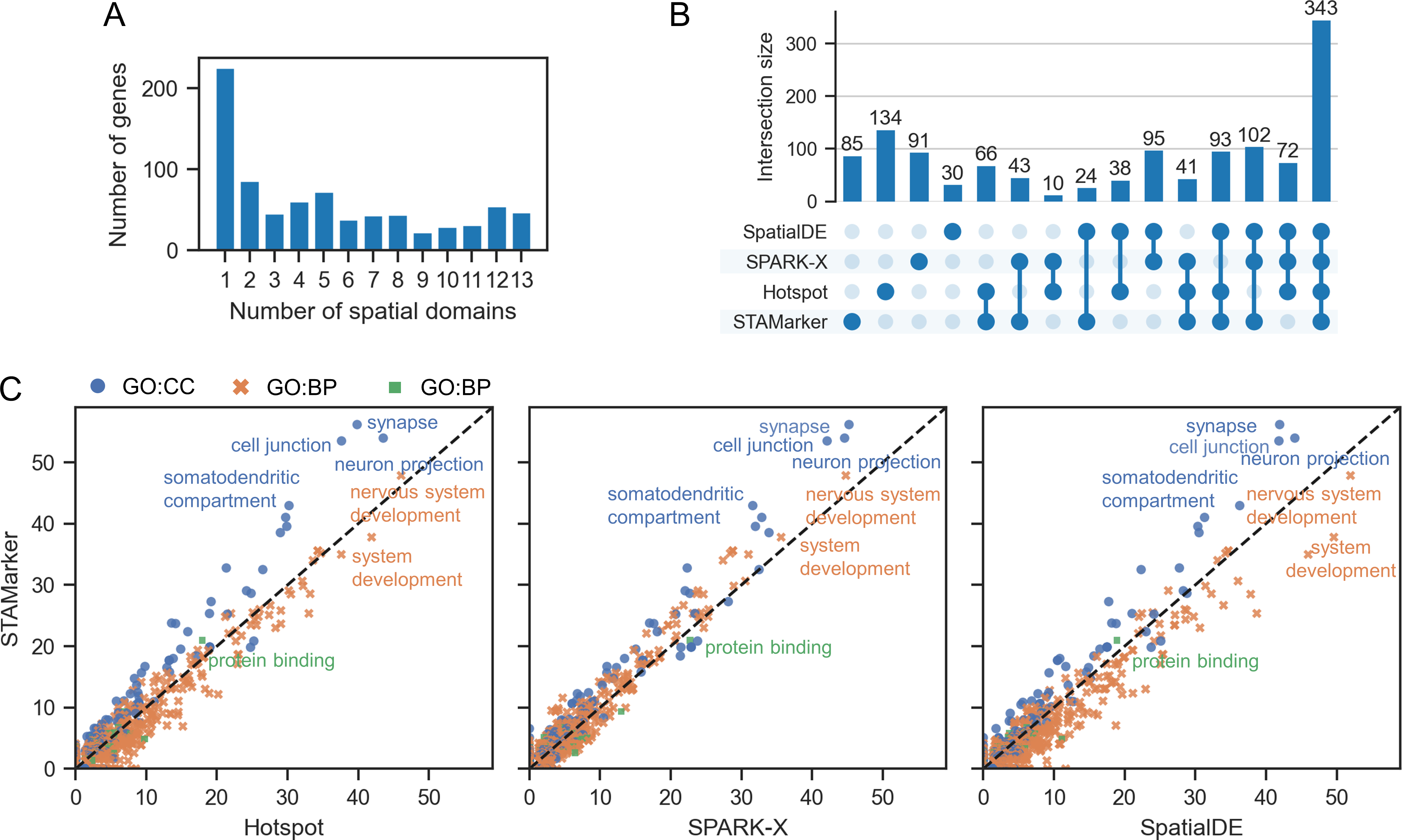

**Figure S2. Additional results on the mouse hippocampus dataset. A**, Histogram of the number of spatial domains to which the SVGs identified by STAMarker belong. **B,** UpSet plot of the numbers of SVGs identified by SpatialDE, SPARK-X, Hotspot, and STAMarker. STAMarker identified 797 SVGs. **D**, Comparison of enrichment significance –log(*p* values) of GO terms enriched with SVGs identified by STAMarker and three compared methods.

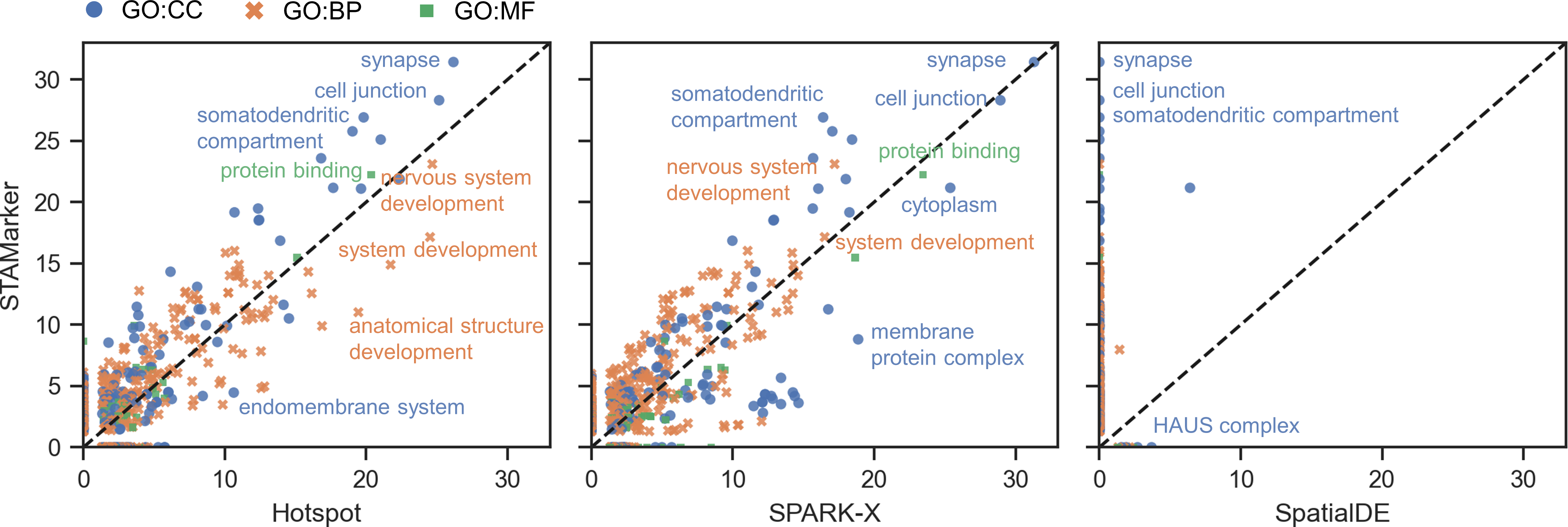

**Figure S3. Additional results on the mouse olfactory bulb dataset.** Comparison of enrichment significance –log(*p* values) of GO terms enriched with SVGs identified by STAMarker and three compared methods.

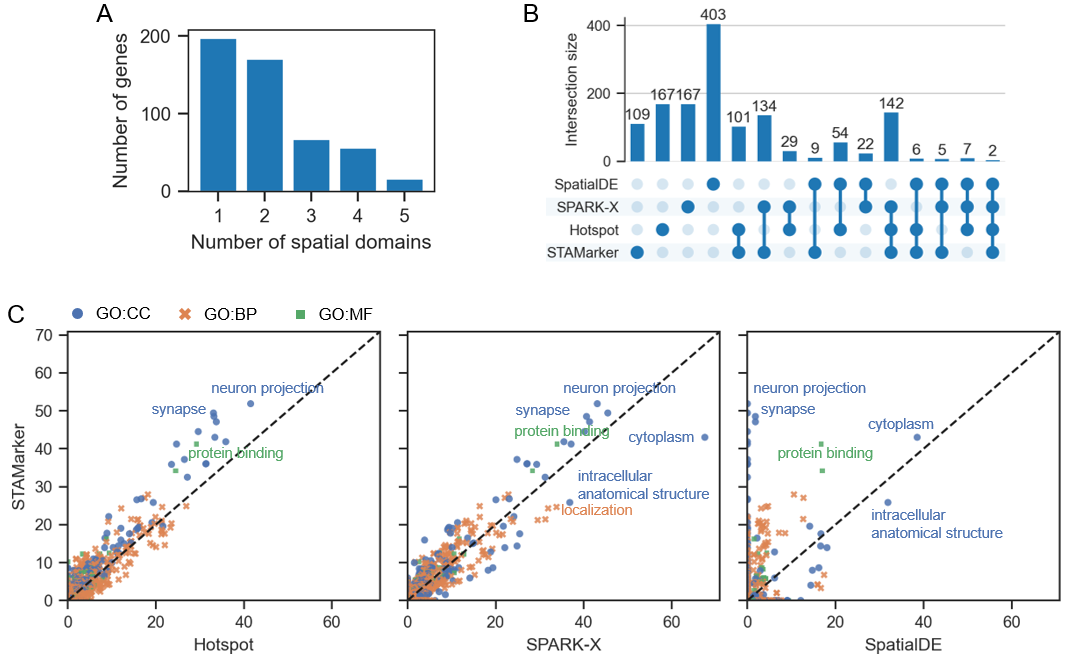

**Figure S4. Additional results on the mouse cerebellum dataset.** **A**, Histogram of the number of spatial domains to which the SVGs identified by STAMarker belong. **B,** UpSet plot of the numbers of SVGs identified by SpatialDE, SPARK-X, Hotspot, and STAMarker. STAMarker identified 437 SVGs. **D**, Comparison of enrichment significance –log(*p* values) of GO terms enriched with SVGs identified by STAMarker and three compared methods.

**Supplementary Tables**

**Table S1.** Description of all ST datasets used in this study.

| Platform | Tissue | Section | #Spots | Related figures | Reference |
| --- | --- | --- | --- | --- | --- |
| 10x Visium | Human dorsolateral prefrontal cortex (DLPFC) | 151507,  151508,  151509,  151510,  151669,  151670,  151671,  151672,  151673,  151674,  151675,  151676 | 4226,  4384,  4789,  4634  3661,  3498,  4110,  4015,  3639,  3673,  3592,  3460 | Figure 2A-E,  Figure S1A-E | [1] |
| Slide-seq V2 | Hippocampus of J20 genetic mouse model with Alzheimer’s disease | SpatialRNA_cropped_  slideseq_j20_rep1 | 15092 | Figure 3A-F,  Figure S2A-C | [2] |
|  | Mouse cerebellum | Cerebellum_Puck_  180819 | 32701 | Figure 5A-G,  Figure S4A-C | [3] |
| Stereo-seq | Mouse olfactory bulb | X | 19109 | Figure 4A-F,  Figure S3 | [4] |
